# Camelot – Intuitive Software for Camera Trap Data Management

**DOI:** 10.1101/203216

**Authors:** Heidi Hendry, Christopher Mann

## Abstract

A variety of software tools for camera trap data management have been produced over the years, though the authors encountered numerous shortcomings with each. Camelot is a new open source cross-platform software application for managing camera trap survey data.

Camelot is designed to be both powerful and easy-to-use. It provides a number of innovative features, such as an extensible reporting system that serves as an integration point with specialised analysis tools. This combination of features and usability makes it a compelling alternative to existing camera trap data management software.

## 1 Introduction

In the field of conservation, it is a common practice to install trail cameras in animal habitats with the express purpose of identifying the occupancy, abundance or behaviour of specific animal species. Trail cameras are placed in outdoor areas that can be inaccessible, depending on weather and other conditions (Fegraus et al., 2011), and often left in the field for months to take pictures with motion sensing triggering photographs of passing animals.

With decreasing cost of suitable trail cameras, camera trap surveys are becoming more prevalent. However, camera trap software has not developed at the same rate. Cataloguing and analysis of camera trap surveys is time-consuming and labor intensive, with available software not always well documented in its use or the types of analyses which will be available.

Camelot is a new open source software tool for managing the data associated with camera trap surveys, specifically for applications in wildlife conservation research. Camelot is designed to be the first step in camera trap survey image classification, and provide versatile outputs that can be used in other software. The overarching goal of Camelot is to provide a modern and intuitive software application for classifying large volumes of camera trap data efficiently and accurately.

## 2 Background

Harris, Thompson, Childs, and Sanderson (2010) developed a camera trap image methodology that many scientists still use today. This methodology involves renaming image files with their timestamp and then sorting the image files into a single-root hierarchical folder structure of the form seen in Figure 1.

**Figure 1.**
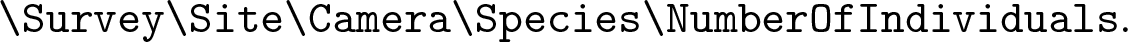
Directory hierarchy for Sanderson & Harris

The researcher subsequently runs the Sanderson & Harris programs to get a report output. The Sanderson & Harris programs are written in FORTRAN (Harris et al., 2010) and are specific to the Microsoft Windows operating system.

Harris et al. program instructions set out a strict sequence of multiple programs in order to transform and analyse the data. Though this is a widely used methodology, many scientists have found this method cumbersome, and problematic. Fegraus et al. (2011) states “Capturing the necessary image meta-data that is required for scientific analyses is particularly challenging”.

Reports from the Harris et al. programs are text files, but are not in a common standard such as tab-delimited or comma-separated-value text files. As a result, these non-standard output formats restrict interoperability with other analysis tools. Additionally, as the images are stored in a single-root hierarchical manner, the reports are survey specific and are not able to produce cross-survey insights.

Furthermore, the approach taken reduces the availability of data validation, allowing simple errors made during “drag & drop” and data entry to result in inconsistencies in any subsequent analysis. The folder hierarchy is fundamentally a denormalised data structure and as such is more prone to data integrity error than a normalised database (Codd, 1970). Whitman and Mattord (2011) states that if users cannot verify the integrity of the data, then the information is of no value. These shortcomings may have contributed to the recent increase in camera trap software development, e.g., TEAM Desktop (Fegraus et al., 2011), Snoopy (Smedley & Terdal, 2014), TRAPPER (Bubnicki, Churski, & Kuijper, 2016). As the technological landscape changes, it is important the tools also evolve to meet the needs and expectations of their users, including supporting a range of operating systems and allowing use by multiple simultaneous people.

Efforts have been made within the conservation community to produce database software in order to address some of the above issues. Two recently introduced projects in the field are TRAPPER, which demonstrates a similar set of design choices to Camelot, and Snoopy. TRAPPER is a database-backed multi-user web applications to be deployed within a Local Area Network. However, unlike Camelot, it requires a standalone database server which necessitates a specialised skillset to install (Bubnicki et al., 2016). Snoopy (Smedley & Terdal, 2014) similarly requires a standalone database server, but is currently still in beta testing. Our experience with Snoopy indicated some problems with operation on a Windows 10 device.

## 3 Camelot

We will now introduce Camelot, a new solution to camera trap data management designed to strike a balance between functionality and usability. The following sections will discuss the key design decisions and features available in Camelot which make it a compelling alternative to existing software applications.

### 3.1 Cross-platform

The architecture of Camelot-a web application which runs on the Java Virtual Machine – makes it highly portable. Its software requirements are the Java Runtime Environment be available on the host machine, and a modern web browser be available on any device accessing Camelot. Camelot has been tested on various versions of Microsoft Windows, macOS and Linux, and on the latest versions of Internet Explorer, Edge, Firefox, Chrome and Safari. Camelot was designed with a responsive user interface and can also be accessed from mobile devices, such as phones and tablets.

### 3.2 Simplified installation and configuration

Software products often rely on an external database, which can be difficult for end users to install and configure, as this oft-times requires specialist knowledge. Camelot avoids the need for a separate database server through the use of an embedded Apache Derby database. While Apache Derby is not as performant or fully-featured as a standalone Database Management System such as PostgreSQL, Camelot has been designed to account for these limitations and has been load tested with datasets of up to 2 million photos.

Once Camelot is downloaded and extracted from its compressed file, there are operating-specific scripts to make starting Camelot straight-forward to use as a desktop application. Those with more advanced technical skills may use the Java archive directly, which provides more fine-grained control around the application’s behaviours. This can be desirable when running Camelot on a server with numerous clients.

Additional configuration is not generally required, though the ability to manage environment-specific configuration, such the TCP port and data directory, is available.

### 3.3 Simplified image import

Camelot’s surveys, sites and cameras are set up through an intuitive web browser user interface. Sites and cameras can also be re-used across multiple surveys which enables subsequent longitudinal analysis. The image import process is a “drag & drop” process, which will also perform validation of the imported data in order to maintain data integrity and guard against human error.

Image import processing involves collecting the available information from the images’ metadata and storing that for reporting. Metadata is extracted from the images using Drew Noakes’ Metadata Extractor library. Camelot provides a workflow that caters for camera trap surveys currently in the field, where, as images are collected, they can be processed immediately and ongoing reports produced. The same workflow can be used to import data for surveys which have been conducted, but are waiting subsequent analysis.

### 3.4 Efficient species identification

Camelot assists with the accurate and efficient entry of taxonomic data using the online Catalogue of Life species database (Roskov et al., 2016). If Camelot is running on an internet-connected computer, then a species lookup for the correct taxonomic description of a species is available, which reduces input time and risk of data entry errors. The species can also be manually entered, which can be useful in the event a species is not in the database, or an internet connection is not available at the time.

Camelot provides a specialised image identification interface called the ‘library’. Here, images can be single-or multi-selected and details of the sighted species identified. The process has been optimised to minimise the amount of human interaction required for each identification in order to further reduce the identification time required. Clicking on an image opens a full resolution version of the image, where the web browser’s zoom features can be used if required. Additional, user-defined input fields can be added to the identification process if required.

### 3.5 User-defined reports

The ability to integrate with other systems used in wildlife conservation research is a core component of Camelot’s role in providing management of camera trap data: the data is able to be exported to suit any number of existing or yet-to-be-created specialised software applications in order to subsequently analyse that data.

Camelot’s primary form of output is a report. A report is a tabular subset of Camelot’s database, provided as a CSV. A report in Camelot is somewhat more sophisticated than a simple export of data though, with the ability to derive calculated columns from existing data, as well as filter, transform, aggregate and sort on that data. For advanced use cases, an arbitrary function may also be supplied in order to process the data.

**Figure 2.**
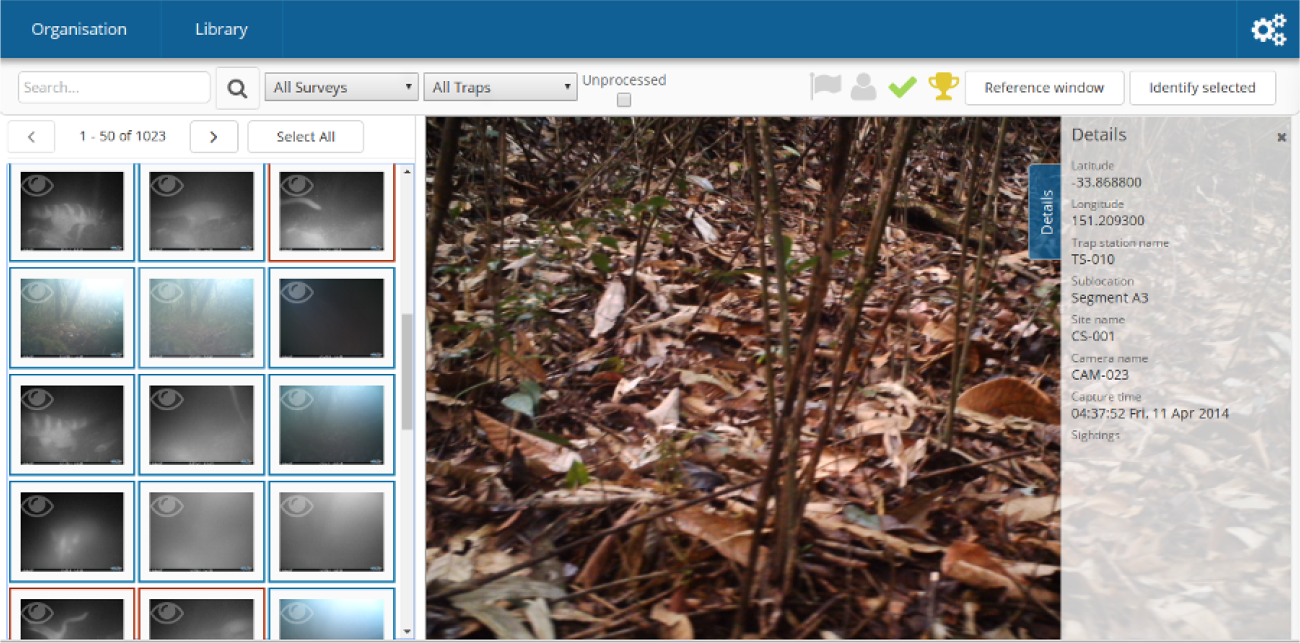
The library is an optimised media management interface. Components of this interface shown here are as follows. **Top:** search bar. Provides searching across all media using quick-access filters and filtering expressions with auto-complete. Also provides a number of status flags which can be assigned to selected media. **Left:** the media collection panel, allows selecting of multiple images simultaneously, pagination of images for performance and manageability, thumbnail borders indicating media status and keyboard navigation. **Centre:** the currently viewed image. **Right:** details panel, a sliding and semi-transparent panel providing details for the currently selected image, including any previous identifications made, and the ability to remove those identifications.

Using these techniques, Camelot provides a number of purpose-specific reports built-in, and is also open to extension by the user. CSV is the output format chosen due to its prevalence and its compatibility with spreadsheeting software enables ad-hoc data exploration and analysis techniques. Several purpose-specific built-in reports are provided to enable subsequent analysis of the data with PRESENCE and the R package camtrapR and any other software utilising similar data formats.

Aside from exposing the data via built-in reports, the reporting functionality is designed to satisfy a number of additional goals:

1. A user should be able to define filters when generating the report. The constraints filtered on will need to be able to vary by report. These should be intuitive to set. This allows data to be restricted during generation, which is particularly important when aggregation occurs and as such could not be subsequently filtered by the user.
2. A user should be able to define their own reports in order to suit new or previously unforeseen use-cases.
3. Consistency in the metrics provided across built-in reports and derivative custom reports should be ensured.

The solution to achieve these goals was to introduce a Domain Specific Language (DSL) which allows reports to be expressed succinctly and declaratively. The DSL is *embedded*, which is to say, that it is evaluated by runtime of Camelot’s implementation language, *Clojure*. Clojure is a general-purpose programming language, and the approach of using an embedded DSL allows users to take full advantage of its capabilities. Dynamic code evaluation facilities make adding a new report, or updating an existing report, a matter of simply saving a file with the report definition to an appropriate configuration folder.

The DSL allows fine-grained control over the generation of reports. The following aspects are exposed in this way, with minimal programming required:

- The columns in the report output,
- The title of the columns,
- The columns over which data is aggregated, where each column has a natural aggregation function built-in,
- Filtering of the data on certain requirements,
- The ordering of results,
- The options presented in the user interface to generate the report, and the title and description shown for the report in the menu,
- The filename of the generated report.

While these are the key areas of functionality the DSL provides, it also exposes advanced functionality whereby report authors can define arbitrarily sophisticated models using the full functionality of the Clojure language. Under this mode of operation, the only constraint imposed by Camelot’s reporting feature that the data be returned in a tabular format, so that it may then be produced as a downloadable CSV. This technique is used in Camelot internally to produce the report to provide interoperability with PRESENCE. Indeed, any report built-in to Camelot is produced by the same DSL available for its users to define their own reports.

### 3.6 Bulk import

In a similar vein to how Camelot’s reporting functions create interoperability with existing analysis tooling by tailoring Camelot’s output to the needs of various other software and processes, the Bulk Import functionality attempts to tailor Camelot’s input to the capabilities of other software. It achieves this by combining three key concepts:

> **Use the most common data interchange format** The ubiquitous nature of spreadsheet software comes due to the ease of data storage and manipulation it offers. These same properties make it more likely that the original data is either stored in, or in a system capable of exporting to, a CSV.
>
> **Systematically discover as much data as possible** Camera trap hardware may store the data available to them within image metadata tags. While there are multiple competing standards for the way in which this data is stored, it does provide a workable mechanism for recording labelled data. Additionally, some methodologies require the structuring of images from camera traps within a directory hierarchy for semantics (Harris et al., 2010). This directory structure is tool-specific and not sufficiently standardised to be used as a basis for import, however components from the directory hierarchy should be available during the import process.
>
> **Provide association between the storage formats** Columns provided by the CSV should be exposed via an interface allowing them to be associated with tables and columns within the database. The user should not need to be aware of the database schema and extensive validation should be performed in real time. The interface should also intelligently select default field associations.

The workflow provided by Camelot to create this highly generalised import is as follows:

1. Specify the directory within which all data for a survey exists.
2. Camelot scans the directory hierarchy, extracts the described data, and produces it as a CSV.
3. The user will review and amend the CSV with any additional data, which may be exported from other software or systems of record.
4. The CSV will be uploaded to Camelot, whereby labelled data from the CSV may be associated with fields in Camelot. At this point Camelot will ensure that the data is valid for the field it is assigned to. For example, if a field must always have a value each record which would be imported must have a value for that field. Similarly, if the field must be a date, it is not valid to import a filename.
5. Once the individual associations are valid, additional validation is performed across the relationships between the records. For example, detecting if a single camera is used in two trap stations at the same time. Should any problem be detected, Camelot will abort the import and notify the user of the problem, including, if possible, the line number of the CSV where the problem was found.
6. Lastly, upon successful validation, the import of the data will commence.

Depending on the original means by which the data was stored, it can be a time consuming task to correct inconsistencies in the CSV for import. The bulk import feature uses the two-step approach to validation in order to detect and notify the user of problems as early as possible. The user may amend and upload a new CSV at any time and Camelot will, insofar as is possible, preserve the existing configuration of the field associations and re-perform the validation.

### 3.7 Data validation

Camelot was designed with the goal of catching data errors and preventing them from tarnishing the integrity of the collected data. Oft-times camera trap images inherently contain errors, such as an incorrect timestamp due to camera malfunction or incorrect configuration. By constraining data input and validating entities as they are added to, or imported into, Camelot, it’s possible to quickly catch and highlight data inconsistencies such as an incorrect timestamp on imported media, session date overlaps and overlapping use of a camera. Furthermore, by encouraging the user to pick their expected species list using the Catalogue of Life database as a guide for species naming, Camelot improves consistency in naming.

For flexibility, Camelot does allow a means to opt-out of many of the constraints imposed on the data to ensure its integrity, however doing so requires it be a deliberate choice by the user.

### 3.8 Longitudinal data collection and reporting

Camelot has a multi-root hierarchical database structure. The single-root hierarchical methodology is not an essential requirement for camera trap software, though most camera trap software has been built with that methodology in mind, Camelot also allows for reporting which is based on sites, species or cameras and is not restricted to a single survey.

Longitudinal studies are more easily facilitated with a multi-root hierarchical structure. Field schools, where surveys are performed at the same site over multiple time periods, may find Camelot a versatile tool to aid in longitudinal reporting.

**Figure 3.**
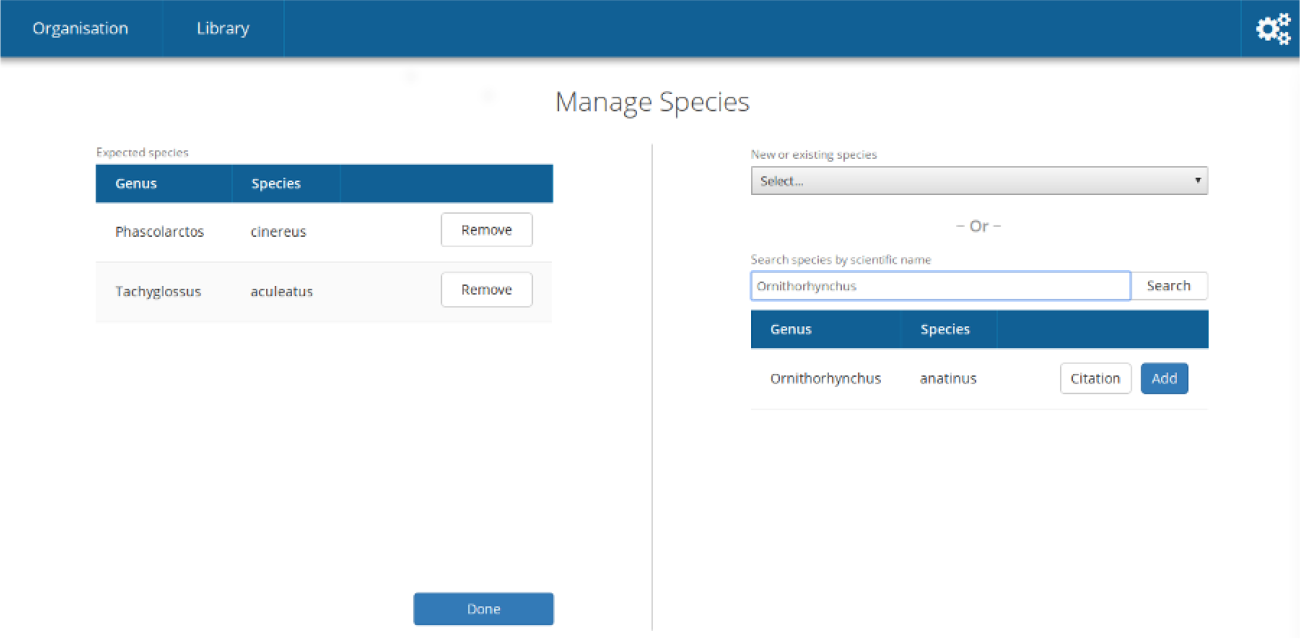
Camelot provides a variety of methods to add species to a survey, including a search interface to the databases provided by Catalogue of Life, and a drop-down menu for species found in other surveys within Camelot.

### 3.9 Open source

Camelot’s source code is available on GitLab and available under the terms of the Eclipse Public License 1.0 or later. Those looking to extend and improve Camelot are encouraged to do so, and can find a contribution guide at the project website.

## 4 Software availability

The Camelot application, source code and documentation are available from https://gitlab.com/camelot-project/camelot.

The Camelot community for questions, support and discussion is located at the Google Group: https://groups.google.com/forum/#!forum/camelot-project.

## 5 Future development

While Camelot has progressed the capabilities of camera trap management software in a number of areas, there is still much room for improvement. The Bulk Import functionality combined with the Survey Export report in Camelot provides its users with some degree of capability towards data sharing. However, while the import requires both the CSV of the data and the media, the Survey Export provides only the CSV; exporting the corresponding media is not facilitated. This impedes data sharing. Furthermore, when received, the data in the CSV must then be massaged to match the path names on the host which will be importing the data.

A future development to Camelot would be extend this functionality so that data can be exported and imported by other users with a minimum of effort. While technologically feasible, it will require some research into the different intents behind instances of data sharing, whether compatibility with other systems and practices is achievable, and, due to the vast amounts of data collected in research conducted with aid of camera traps, the most suitable means to make-available the data to be shared.

Meanwhile, Camelot’s primary output format lowers the barrier to introducing custom analyses into a camera trap application directly. However defining custom reports in Camelot still requires significant technical ability. The DSLs serve not as the end goal but as an intermediate step towards providing for user defined reports via the web interface alone. Currently, implementing some types of reports may require knowledge of the report generation internals. In the future the authors hope to extend the current facilities for report generation and in doing so devise a simple web interface for creating custom reports without requiring the user to have any knowledge of the DSL or internals of Camelot’s report generation process.

## 6 Conclusion

Camelot is offered to the conservation science community as a species identification and reporting tool. Camelot is a flexible and simple cross-platform software that speeds up the analysis of camera trap survey images.

Camelot accommodates both the single researcher, as well as teams of researchers. The single researcher should find Camelot easy to install and configure, and should be straightforward to install on their own computer. Small-and medium-sized teams may benefit from having Camelot installed and available on a workstation or server, which any number of them can connect to and access concurrently.

The reporting functionality of Camelot provides valuable output for analysis, combining compatibility with existing specialised tools with a useful output format for more general tools such as spreadsheet applications for data exploration and simple analyses. Reporting modules lower the programming knowledge necessary for users to produce their own analyses directly from camera trap software.

With quick image import and species identification, Camelot will reduce the processing time burden on scientists, allowing them to begin the analysis sooner. Camelot’s pre-defined reports are already compatible with existing conservation science software, and Camelot’s flexibility for creating customised reports ensures that it will be relevant in the future. Furthermore, Camelot’s multi-root hierarchy allows for different analyses on existing data, such as site-based, species-based or even camera-based. We believe that Camelot is a valuable new tool for camera trap data management.

## 7 Acknowledgements

Input into design was provided by Dr Benjamin Rawson of Fauna & Flora International - Vietnam. With thanks to the Fauna & Flora International camera trappers in Myanmar and Indonesia, especially Wido Albert, Grant Cornette, and Patrick Oswald for detailed feedback about usability and preferred report outputs. Further thanks to all other Fauna & Flora International Camelot Beta Testers for their support and feedback.

## References

Bubnicki, J. W., Churski, M., & Kuijper, D. P. (2016). Trapper: An open source web-based application to manage camera trapping projects. Methods in Ecology and Evolution, 7(10), 1209–1216.

Codd, E. F. (1970). A relational model of data for large shared data banks. Communications of the ACM, 13(6), 377–387.

Fegraus, E. H., Lin, K., Ahumada, J. A., Baru, C., Chandra, S., & Youn, C. (2011). Data acquisition and management software for camera trap data: A case study from the team network. Ecological Informatics, 6(6), 345–353.

Harris, G., Thompson, R., Childs, J. L., & Sanderson, J. G. (2010). Automatic storage and analysis of camera trap data. The Bulletin of the Ecological Society of America, 91(3), 352–360.

Roskov, Y., Abucay, L., Orrell, T., Nicolson, D., Flann, C., Bailly, N., … Wever, A. D. (2016). http://www.catalogueoflife.org/annual-checklist/2016. Species 2000 & ITIS Catalogue of Life, 2016 Annual Checklist.

Smedley, R. & Terdal, E. (2014). Snoopy: Portable software for capture-recapture surveys.

Whitman, M. E. & Mattord, H. J. (2011). Principles of information security. Cengage Learning.

